# Meiosis leads to pervasive segregation distortion and copy-number variation in accessory chromosomes of the wheat pathogen *Zymoseptoria tritici*

**DOI:** 10.1101/214882

**Authors:** Simone Fouché, Clémence Plissonneau, Bruce A. McDonald, Daniel Croll

## Abstract

Meiosis is one of the most conserved molecular processes in eukaryotes. The fidelity of pairing and segregation of homologous chromosomes has a major impact on the proper transmission of genetic information. Aberrant chromosomal transmission can have major phenotypic consequences, yet the mechanisms are poorly understood. Fungi are excellent models to investigate processes of chromosomal transmission, because many species have highly polymorphic genomes that include accessory chromosomes. Inheritance of accessory chromosomes is often unstable and chromosomal losses have little impact on fitness. We analyzed chromosomal inheritance in 477 progeny coming from two crosses of the fungal wheat pathogen *Zymoseptoria tritici*. For this, we developed a high-throughput screening method based on restriction site associated DNA sequencing (RAD-seq) that generated dense coverage of genetic markers along each chromosome. We identified rare instances of chromosomal duplications (disomy) in core chromosomes. Accessory chromosomes showed high overall frequencies of disomy. Chromosomal rearrangements were found exclusively on accessory chromosomes and were more frequent than disomy. Accessory chromosomes present in only one of the parents in an analyzed cross were inherited at significantly higher rates than the expected 1:1 segregation ratio. Both the chromosome and the parental background had significant impacts on the rates of disomy, losses, rearrangements and segregation distortion. We found that chromosomes with higher sequence similarity and lower repeat content were inherited more faithfully. The large number of rearranged progeny chromosomes identified in this species will enable detailed analyses of the mechanisms underlying chromosomal rearrangement.

## Introduction

Sexual reproduction requires chromosomes to undergo meiosis, whereby homologous chromosomes pair, recombine and finally separate and migrate to opposite poles of the meiotic cell. Meiosis is a highly conserved process initiated by the pairing of homologous chromosomes that first recognize one another and then establish recombination-dependent links between homologs to form the synaptonemal complex (reviewed in Roeder, 1997). This is followed by two divisions, first to separate homologous chromosomes and then to separate sister chromatids. While accurate pairing of homologs is essential for the faithful segregation of chromosomes (Naranjo, 2012), chromosomes can pair along their entire length or in a segment-specific manner where only some regions align (Roeder, 1997). This suggests that the length and degree of sequence similarity can affect homolog identification and pairing. After pairing, recombination produces crossovers that physically link homologs, mediate proper segregation, and thereby preserves chromosomal integrity (Baker et al., 1976; Hassold and Hunt, 2001; Mather, 1938). Recombination between misaligned repetitive sequences can generate length variation among the daughter chromosomes (Montgomery et al., 1991). After pairing and recombination, segregation occurs via centromeres that bind to chromosome proteins and mediate accurate segregation to the opposite poles of the cell.

Aberrant transmission of chromosomes from one generation to the next, including partial and whole chromosome duplications or losses, are caused largely by erroneous pairing during meiosis. Such duplication and loss events can affect a large number of genes and alter gene expression across the genome (Harewood and Fraser, 2014). The most dramatic copy number variation is aneuploidy. Unequal sets of chromosomes result from non-disjunction and are the leading genetic cause of miscarriages in humans (Hassold and Hunt, 2001). Atypical phenotypes associated with aneuploid states are caused by gene dosage imbalances that can cause severe defects (Torres et al., 2008). In general, aneuploidy and chromosomal rearrangements are associated with lower fitness (Torres et al., 2008), but in rare circumstances, errors during meiosis can provide adaptive genetic variation. For example, in the human pathogenic fungi *Cryptococcus neoformans* and *Candida albicans*, specific aneuploidies contribute to drug resistance (Ngamskulrungroj et al., 2012; Selmecki et al., 2006, 2008; Sionov et al., 2010). Adaptive aneuploidy is frequently associated with response to stressful environments (Chen et al., 2012). The dosage imbalance and altered stoichiometry due to additional copies of genes on a duplicated chromosome may not be beneficial under normal conditions, but can become beneficial under stress (Pavelka et al., 2010a, 2010b). In pathogenic fungi, aneuploidy often occurs for only a restricted number of chromosomes, however the mechanisms determining the rate of aneuploidy generation and its maintenance are poorly understood.

Aneuploidy also plays an important role in several plant pathogenic fungi. Several important plant pathogens have highly dynamic genomes with chromosomes that show significant length and number polymorphisms within the species. This chromosomal plasticity is often restricted to a well-defined set of accessory chromosomes. This bipartite genome structure, characterized by an accessory genome region that is rapidly diversifying and a core genome region that remains conserved, can be associated with the trajectory of pathogen evolution (Croll and McDonald, 2012; Dong et al., 2015). The accessory region is often rich in transposable elements that drive chromosomal rearrangements, deletions and duplications (Zhang et al., 2011). Accessory chromosomes are not shared among all members of a species, therefore these chromosomes can contribute significantly to polymorphism within a species. Importantly, many plant pathogens have been shown to harbor pathogenicity loci on accessory chromosomes (Möller and Stukenbrock, 2017). In contrast, the core regions encode essential functions required for survival and reproduction. Plant pathogenic fungi provide particularly powerful models to investigate factors affecting the transmission of chromosomes through meiosis because of their extreme chromosomal plasticity, the ubiquity of sexual reproduction and their experimental tractability.

The fungal wheat pathogen *Zymoseptoria tritici* provides a striking example of genome plasticity. The bipartite genome consists of 13 core and up to eight accessory chromosomes that exhibit significant length polymorphism within and among field populations (Croll and McDonald, 2012; Goodwin et al., 2011). Chromosomal rearrangements played an important role in adaptation to different host genotypes (Hartmann et al., 2017). The accessory chromosomes are highly unstable through meiosis and were shown to undergo rearrangements, segregation distortion and non-disjunction (Croll et al., 2013; Wittenberg et al., 2009). Furthermore, the pathogen tolerates aneuploidy and chromosomal rearrangements in the core and accessory genomes (Croll et al., 2013; Schotanus et al., 2015; Wittenberg et al., 2009). Hence, this species is an ideal model to analyze patterns of aberrant chromosomal transmission.

In this study, we analyze the mechanisms that affect the fidelity of chromosomal inheritance through meiosis, including identification of chromosomal rearrangements, losses and duplications. For this, we screened hundreds of progeny genotypes generated from two independent crosses and determined the rate of aneuploidy, patterns of rearrangement and distortions in transmission rates. Finally, we investigated whether factors such as length similarity, synteny, recombination rate and repetitive element content affected the fidelity of chromosomal inheritance.

## Materials and methods

### Generation of sexual crosses

Two crosses were performed between four parental *Z. tritici* isolates collected from two Swiss wheat fields separated by approximately 10 km. Isolate ST99CH3D1 was crossed with isolate ST99CH3D7 (hereafter abbreviated 3D1 and 3D7) and isolate ST99CH1A5 was crossed with isolate ST99CH1E4 (abbreviated 1A5 and 1E4), producing 359 and 341 progeny, respectively. The genomes of all four parental isolates were sequenced using Illumina technology (Torriani et al., 2011) and are available under the NCBI SRA accession numbers SRS383146 (3D1), SRS383147 (3D7), SRS383142 (1A5) and SRS383143 (1E4). The parental isolates were already genetically characterized and have been phenotyped for virulence and many other traits (Croll et al., 2013; Zhan et al., 2005). Full sib families were produced by coinfecting wheat leaves with parental strains using the crossing protocol described by Kema et al. (1996). Potential clones were identified as offspring sharing >90% SNP identity. Only one randomly selected isolate from any group of potential clones was kept for further analysis, reducing the number of unique offspring to 263 in the 3D1 × 3D7 cross and to 261 in the 1A5 × 1E4 cross (Lendenmann et al., 2014).

### Reference alignment using restriction-associated DNA sequencing (RAD-seq)

We used RAD-seq (Baird et al., 2008) for large-scale sequence genotyping as described previously (Croll et al., 2015). Briefly, the RAD-seq protocol (Etter et al., 2011) was applied to *Z. tritici* by using the *Pst*I restriction enzyme to digest 1.3 μg of DNA extracted with the DNAeasy plant mini kit (QIAGEN Inc., Basel, Switzerland) for each offspring. After digestion and adapter annealing, the pooled DNA was sequenced on an Illumina HiSeq2000 using a paired-end 100 bp library. Pools contained ~132 progeny, six different Illumina TruSeq compatible P2 adapters and 22 P1 adapters with unique barcodes. Progeny DNA with the same P2 adapter were distinguishable by using the unique barcodes ligated to the P1 adapters.

Illumina reads were quality trimmed using Trimmomatic v. 0.30 (Bolger et al., 2014) and separated into distinct sets for each progeny based on the P1 adapter using FASTX toolkit v 0.13 (http://hannonlab.cshl.edu/fastx_toolkit/). Reads were aligned to the gapless telomere to telomere IPO323 reference genome (assembly version MG2, Sept, 2008) (Goodwin et al., 2011) with the short-read aligner version of bowtie 2.1.0 (Langmead and Salzberg, 2012) using the default parameters for sensitive-end-to-end alignment (-D 15; -R 2; L- 22; -I S, 1,1.15). The same parameters from trimming and reference assembly were used to align the four parental genome sequences (Croll et al., 2013) to the reference genome (IPO323). RAD-seq aligned reads are available under the NCBI BioProject accession numbers PRJNA256988 and PRJNA256991.

### Determining chromosome number and length polymorphisms based on coverage

Restriction sites cut by *Pst*I were identified *in silico* using the EMBOSS restrict program (http://emboss.source-forge.net/apps/restrict.html). Thereafter, the coverage of RAD-seq reads mapping to the restriction sites was determined using the BEDtools v. 2.25.0 intersectBed and coverageBed commands (Quinlan and Hall, 2010). Reads were counted if the mapping quality score was ≥20. The coverage of the sequenced parent genomes was determined following the same procedure. Progeny with a median read coverage of <20X were excluded from further analyses to avoid biases introduced by low-coverage data, resulting in fewer isolates being included in this analysis than in previous studies (Lendenmann et al., 2016, 2014, Stewart et al., 2017, 2017). We used normalized read counts to detect chromosomal anomalies, where those with a normalized coverage close to zero (<0.3) were classified as missing, those with a normalized coverage close to one (>=0.7 and <1.3) were classified as present and those with a normalized coverage close to two (>=1.7) were classified as disomic (Figure 1A). Partially deleted and partially duplicated chromosomes were identified based on a normalized coverage ratio of >=0.3 and <0.7 or >1.3 and <1.7, respectively. Deviations from Mendelian inheritance for accessory chromosomes present in only one of the parents were determined using a chi-squared (χ^2^) test.

**Figure 1:**
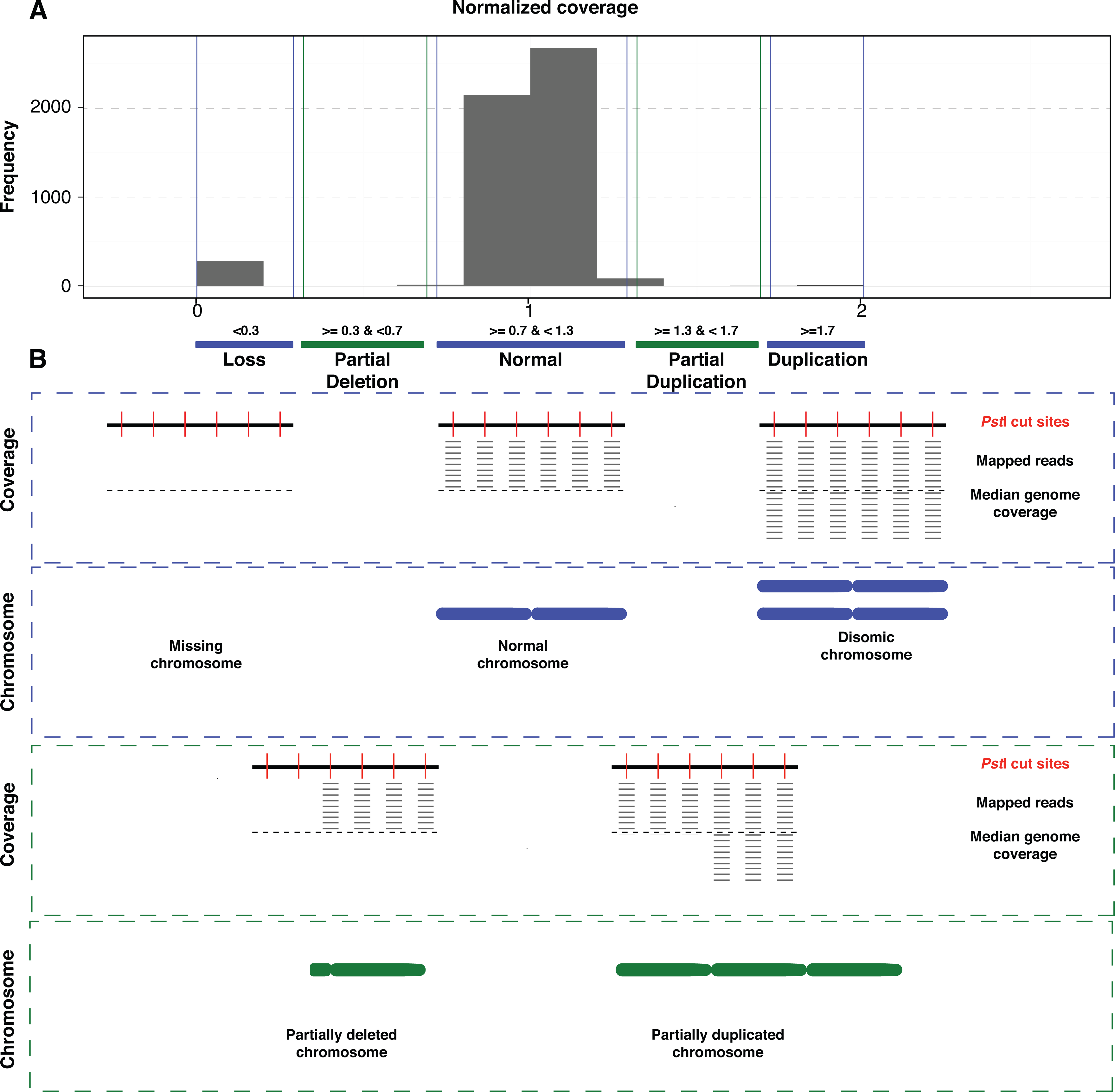
Procedure to detect chromosomal anomalies. A) Reads mapped to the *Pst*I restriction sites were used to analyze coverage across the genome. Sequencing data was generated by restriction site associated DNA sequencing (RAD-seq). The normalized coverage represents the coverage of each chromosome normalized to the median coverage of all chromosomes of the same progeny. The normalized coverage distribution of progeny from cross 3D1x3D7 is shown with the cutoffs used to detect a whole chromosome loss (ratio < 0.3), partial deletion (ratio 0.3-0.7), normal transmission (0.7-1.3), partial duplication (1.3-1.7) and whole chromosome duplication (>1.7). B) Schematic overview of read coverage expected for complete chromosome losses and duplications (in blue). Partial deletions and duplications are shown in green.

### Distinguishing between homozygous and heterozygous disomy

Single-nucleotide polymorphism (SNP) calling was performed using Freebayes (Version 1.0.2_1 1.1.0) (Garrison and Marth, 2012) using the bamfiles of each isolate mapped to the IPO reference genome. We used the parameters no-indels, no-mnps, no-complex and ploidy 2. Then we filtered for sites that differed between the parents (maf 0.2) and considered only these regions to determine whether disomic chromosomes originated from one or both parents. We also filtered for depth (minDP 30) and quality (minQ 30). The VCF tools -het function was used to determine the number of homozygous sites (O(hom)) and the total number of sites. We determined the ratio of homozygous sites to the total number of sites and defined those with a ratio >0.6 as homozygous while those with a ratio <0.4 were defined as heterozygous. All other cases were considered to be ambiguous.

### Chromosome instability and recombination rate, chromosome length, synteny and transposable element content of the parent chromosomes

We correlated chromosome instability with the percentage length difference in homologs among the parents and recombination rates based on the recombination rates reported in Croll et al (2015). We also correlated synteny and the fidelity with which chromosomes were inherited using the NUCmer pipeline from MUMmer (version 3.23) software (Kurtz et al., 2004) to determine the sequence similarity between two homologous chromosomes. The minimum cluster length was set to 50 and we used the –mum option to anchor matches that were unique in both the reference and query sequence. The TEs in the parent genomes were annotated using RepeatMasker (http://www.repeatmasker.org) and the transposable element library compiled for *Z. tritici* and its sister species (Grandaubert et al., 2015). The percentage of TEs on a chromosome was compared to the likelihood of being inherited with high fidelity. We also compared the frequency of disomic chromosomes with the frequency of rearrangements for all chromosomes in both crosses.

### Analyses of progeny phenotypes

Progeny from both crosses were phenotyped for percentage of leaf area covered by lesions (PLACL), pycnidia density (pycnidia/cm^2^ leaf area), pycnidia size (mm^2^) and pycnidia melanization on seedlings of the wheat cultivars Runal and Titlis in a previously described glasshouse-based assay (Stewart et al, 2017). Grey values were previously shown to be a good measure for melanization (Lendenmann et al., 2014). The assay was repeated three times over three consecutive weeks, resulting in three biological replicates and six total replicates per isolate-cultivar pair. Automated image analysis of the second leaf was performed at 23 dpi as previously described (Stewart and McDonald, 2014). Progeny were also phenotyped for temperature sensitivity, growth morphology and fungicide sensitivity (Lendenmann et al., 2015, 2016). Phenotypes were compared in normal progeny and progeny with “abnormal” (partially deleted, partially duplicated, disomic or absent) chromosomes to determine if particular chromosome genotypes were associated with outlier virulence, fungicide resistance, temperature sensitivity or growth rate phenotypes. These analyses were performed in R.

## Results

### Mapping RAD-seq reads to the reference genome

The chromosome state (absent, present or duplicated) was determined for each chromosome of the four haploid parental isolates (3D1, 3D7, 1A5, 1E4) and 477 progeny, using RAD-seq reads generated for each progeny mapped to the IPO323 reference genome. The 3D1 and 1A5 parents had all 21 chromosomes, while the 3D7 and 1E4 parents were missing four and one accessory chromosomes, respectively (Croll et al., 2013). We selected the parental isolate from each cross that carried all 21 chromosomes (3D1 and 1A5) as a reference. We mapped whole-genome sequencing data of the two selected parents against the IPO323 reference genome and identified regions missing in the parental genomes. Missing regions were not expected to show coverage in any of the progeny chromosomes and were excluded from further analyses. For each progeny, we calculated the coverage for each chromosome and compared this to the median coverage of all chromosomes for that isolate (Figure 1). The normalized coverage per chromosome was close to one for the large majority of the chromosomes (Supplementary Figure 1). The mean normalized coverage ratio was 0.96 and 0.95 for the progeny from cross 3D1x3D7 and cross 1A5x1E4, respectively.

### Patterns of chromosome transmission in the two crosses

Analyzing normalized read coverage among progeny revealed high rates of chromosome losses in both crosses. In cross 3D1x3D7, accessory chromosomes 16, 17, 19 and 20 were present in both parents but were missing in 1.6%, 4.4%, 0.4% and 1.2% of the progeny, respectively (Figure 2A). In the 1E4x1A5 cross, accessory chromosomes 14, 15, 16, 18, 19, 20 and 21 were present in both parents but were absent in 7.5%, 2.2%, 4.8%, 6.1%, 2.2%, 1.8% and 4.4% of the progeny, respectively (Figure 2B). We found no progeny lacking a core chromosome in either of the crosses.

**Figure 2:**
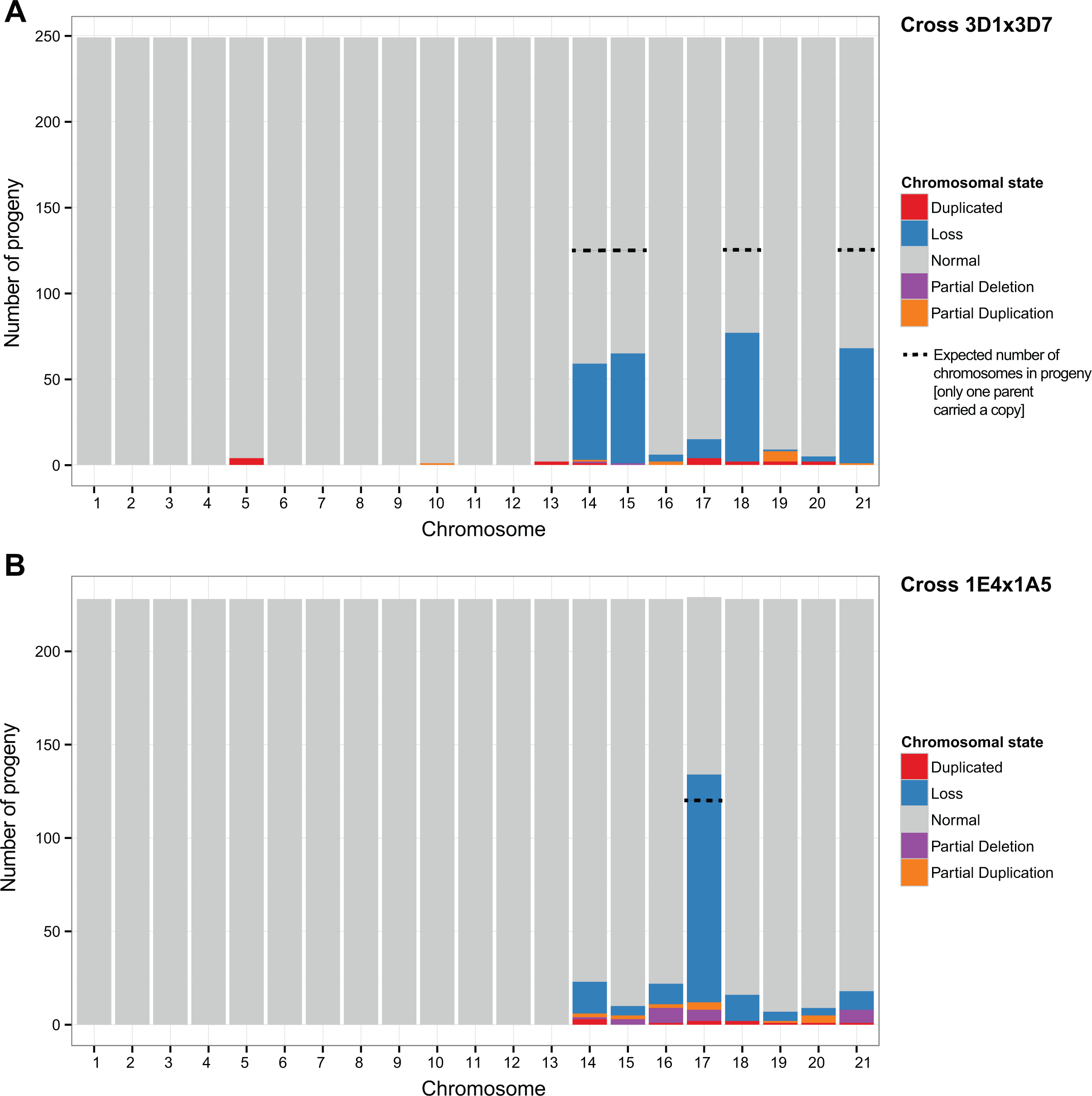
Summary of the total chromosome anomalies in the progeny of two crosses. Normal, disomic, lost and rearranged (partially duplicated or deleted) chromosomes are shown separately for cross 3D1x3D7 and 1E4x1A5. Dotted lines show the expected number of progeny for chromosomes that were present in only one of the two parental isolates.

We also identified numerous instances of disomy in progeny accessory chromosomes. In cross 3D1x3D7, chromosomes 17, 19 and 20 were present in two copies in 1.6%, 0.8% and 0.8% of the progeny, respectively (Figure 2A). Interestingly, 2.4% of the progeny were disomic for a core chromosome, with 1.6% of the progeny disomic for chromosome 5 and 0.8% disomic for chromosome 13. No disomic core chromosomes were identified in cross 1E4x1A5 (Figure 2B), but 1.3% of the progeny were disomic for chromosome 14, 0.9% were disomic for chromosome 18 and chromosomes 16, 19, 20 and 21 were each disomic in 0.4% of the progeny.

Segregation patterns that differed from the expected 1:1 ratio were observed for several chromosomes that were present in only one of the two parents of a cross. In the 3D1x3D7 cross, chromosomes 14, 15, 18 and 21 were absent in the 3D7 parent, hence we expected these chromosomes to be absent in half of the progeny. Instead, chromosomes 14, 15, 18 and 21 were absent in only 22.5%, 25.7%, 30.1% and 26.9% of the progeny, respectively (Figure 2A). The inheritance of these chromosomes are significant departures from the canonical Mendelian ratio (chromosome 14: χ^2^=37.7, *p* < 0.001, chromosome 15: χ^2^=29.4, *p* < 0.001, chromosome 18: χ^2^=19.7, *p* < 0.001 and chromosome 21: χ^2^=26.6, *p* < 0.001). In the 1E4x1A5 cross, chromosome 17 was missing in 53.5% of the progeny and did not exhibit segregation distortion (χ^2^=0.56, *p* = 0.3) (Figure 2B). Disomy was also found for several accessory chromosomes that were present in only one of the parents. In cross 3D1x3D7, additional copies of chromosome 14 and 18 were identified in 0.4% and 0.8% of the progeny, respectively (Figure 2A). In cross 1E4x1A5, chromosome 17 was disomic in 0.9% of the progeny (Figure 2B).

Disomic chromosomes can either be heterozygous, carrying one of each parental chromosomal copy, or homozygous if the disomy arose from a single parental chromosome (Figure 3). To distinguish these scenarios, we analyzed disomic progeny chromosomes and restricted the analyses to cases where both parents were carrying a chromosomal copy. In the 3D1x3D7 cross, 59% (10/17 cases) of the disomic isolates were heterozygous, with a chromosome originating from each parent and 29% (5/17 cases) of the disomic isolates were homozygous, with both chromosomes originating from one parent (Figure 4A). In the case of chromosomes 14 and 18, the chromosomes could only originate from one parent. In cross 1E4x1A5, 5 of the 11 disomic isolates were homozygous, three disomic isolates had chromosomes originating from both parents and the other three cases were ambiguous. As before, chromosome 17 could only have originated from one of the parents.

**Figure 3:**
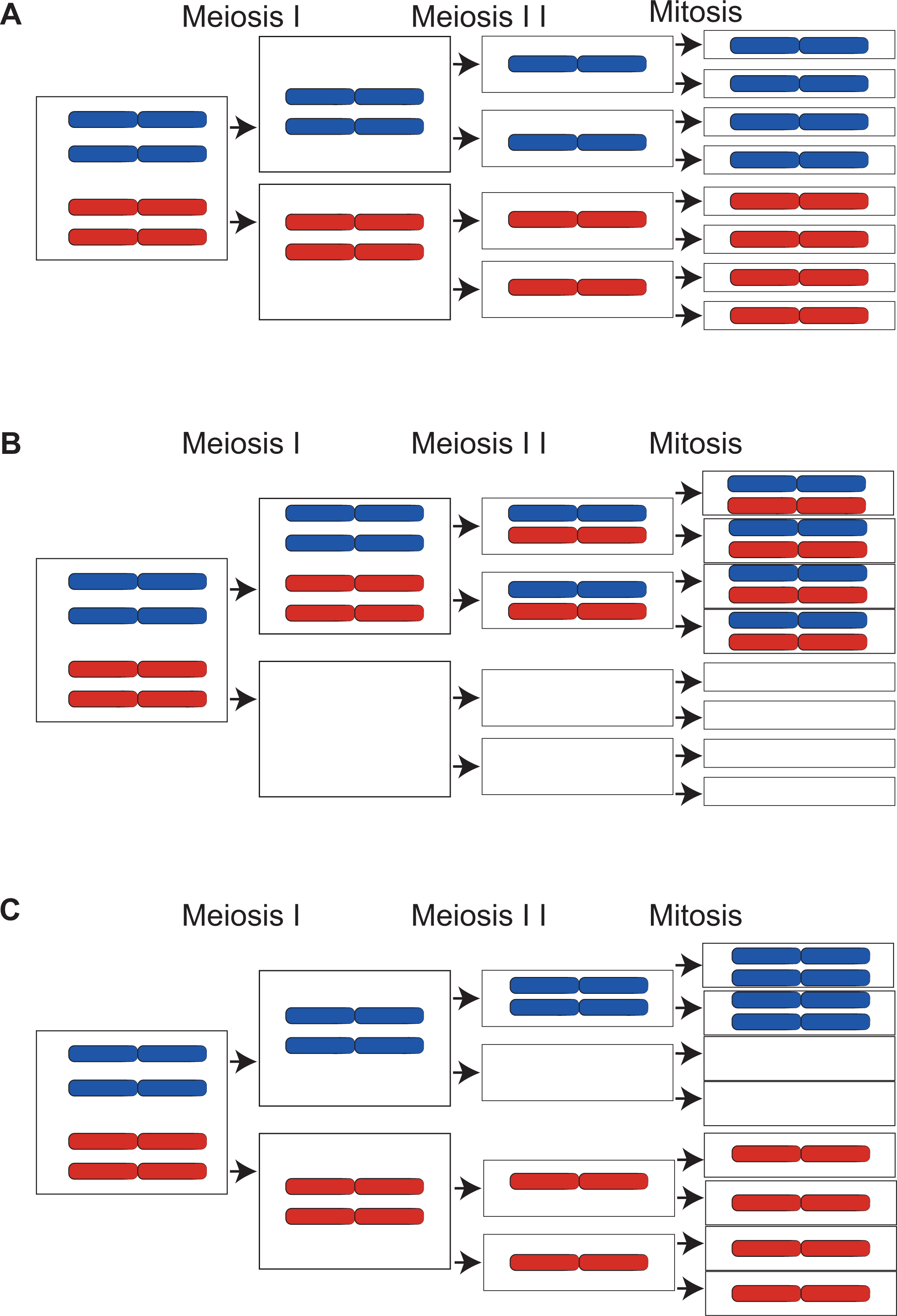
Schematic overview of how chromosomal non-disjunction can result in chromosome loss or disomy. A) During canonical meiosis, the haploid nuclei from the two parents fuse resulting in a single diploid nucleus. Parental chromosomes are shown with distinct colors. Chromosomes go through meiosis I and II, followed by mitosis, resulting in 8 haploid ascospores. Chromosome loss or disomy can occur as a result of homologous chromosomes failing to segregate during meiosis I (B), resulting in heterozygous disomy with one chromosome originating from each of the parents. The alternative is the failure of sister chromatid segregation during meiosis II (C), generating homozygous disomic progeny with both copies of the chromosome originating from the same parent.

**Figure 4:**
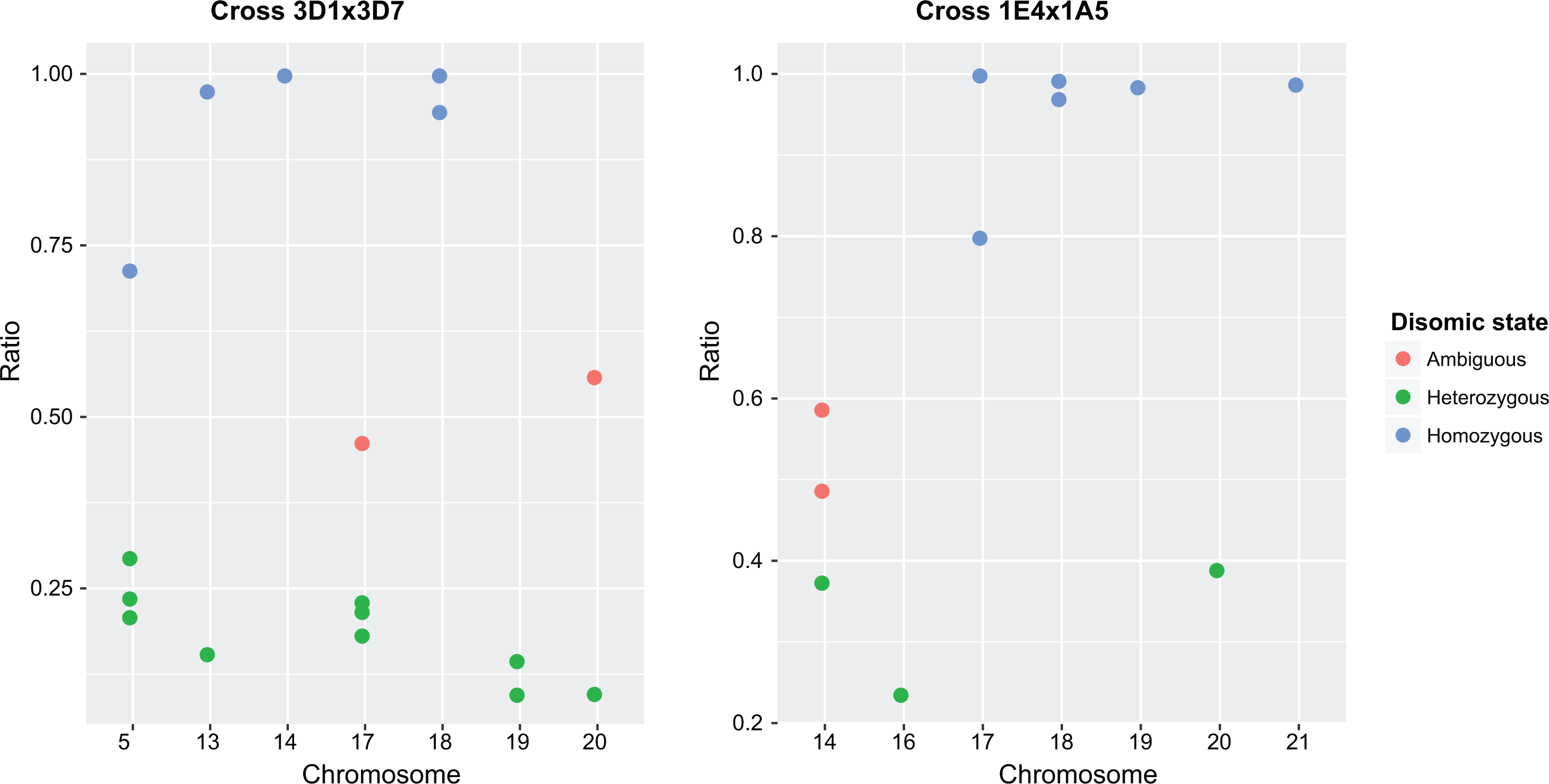
Identification of heterozygous and homozygous disomic chromosomes in cross 3D1x3D7 and cross 1E4x1A5. Single nucleotide polymorphism (SNP) loci were screened on progeny chromosomes that showed evidence for disomy. SNPs were genotyped as either homozygous, containing only one of the parental alleles, or heterozygous if both parental alleles were found. The ratio represents the number of homozygous SNPs compared to the total number of genotyped SNPs. Individual dots represent each of the disomic progeny chromosomes identified in the two crosses. Due to uncertainties in SNP calling, we used cutoffs to assign progeny chromosomal states. Chromosomes with a ratio <0.4 were assigned as heterozygous disomic, likely resulting from nondisjunction at meiosis II, >0.6 as homozygous disomic, likely resulting from nondisjunction at meiosis I, and ratios between 0.4-0.6 were assigned as ambiguous.

### Meiosis generates novel chromosome length polymorphism

In order to identify partially deleted or duplicated chromosomes in the progeny, we investigated chromosomes which had a normalized coverage between 0.3 and 0.7, and between 1.3 and <1.7 (Figure 1). In cross 3D1x3D7 (Figure 2A), partial deletions were identified for chromosomes 14 (0.4% of offspring) and 15 (0.4%). Partial duplications were detected for chromosomes 14 (0.4%), 16 (0.8%), 19 (2.4%) and 21 (0.4%). We also identified one isolate which may have a partially duplicated core chromosome 10. In cross 1E4x1A5 (Figure 2B), partial duplications were detected in the progeny for chromosomes 14 (0.9%), 15 (0.9%), 16 (0.9%), 17 (1.8%), 19 (0.4%) and 20 (1.8%). Partial losses were identified for chromosomes 14 (0.4%), 15 (1.3%), 16 (3.5%), 17 (2.6%) and 21 (3.1%).

We identified some progeny with multiple chromosomal anomalies, however these associations did not significantly deviate from a random expectation. In cross 3D1x3D7, isolate 89.1 was disomic for chromosome 13 and had a large, partial duplication of chromosome 10 while isolate 137.2 had partial duplications of chromosomes 16, 19 and 21. In cross 1E4x1A5, isolate B23.1 was disomic for chromosome 20 and had partial deletions of chromosomes 17 and 21. This isolate also had a partially duplicated chromosome 14. Isolate B24.2 also had partial deletions of chromosomes 17 and 21. Isolate C44.2 had partially deleted chromosomes 16 and 21. Isolate B50.1 was disomic for chromosome 17 and had a partially deleted chromosome 21. Isolate A57.1 was disomic for chromosome 14 and had a partially duplicated chromosome 16.

In cross 3D1x3D7, we found twelve progeny isolates with partial deletions and duplications. Seven of these partial aneuploidies affected chromosomal segments near the telomeric ends (Supplementary Figure 2). Isolate 89.1 had a normalized coverage ratio for chromosome 10 of 1.63 suggesting a partial duplication. However, the coverage along the chromosome was homogeneous with no apparent duplicated chromosomal regions when compared to the parent chromosomes (Supplementary Figure 2). We considered such cases as ambiguous duplications. In cross 1A5x1E4, we found 40 partial deletions and duplications, of which 19 were ambiguous and 15 occurred in chromosomal segments near the telomeric ends (Figure 5).

**Figure 5:**
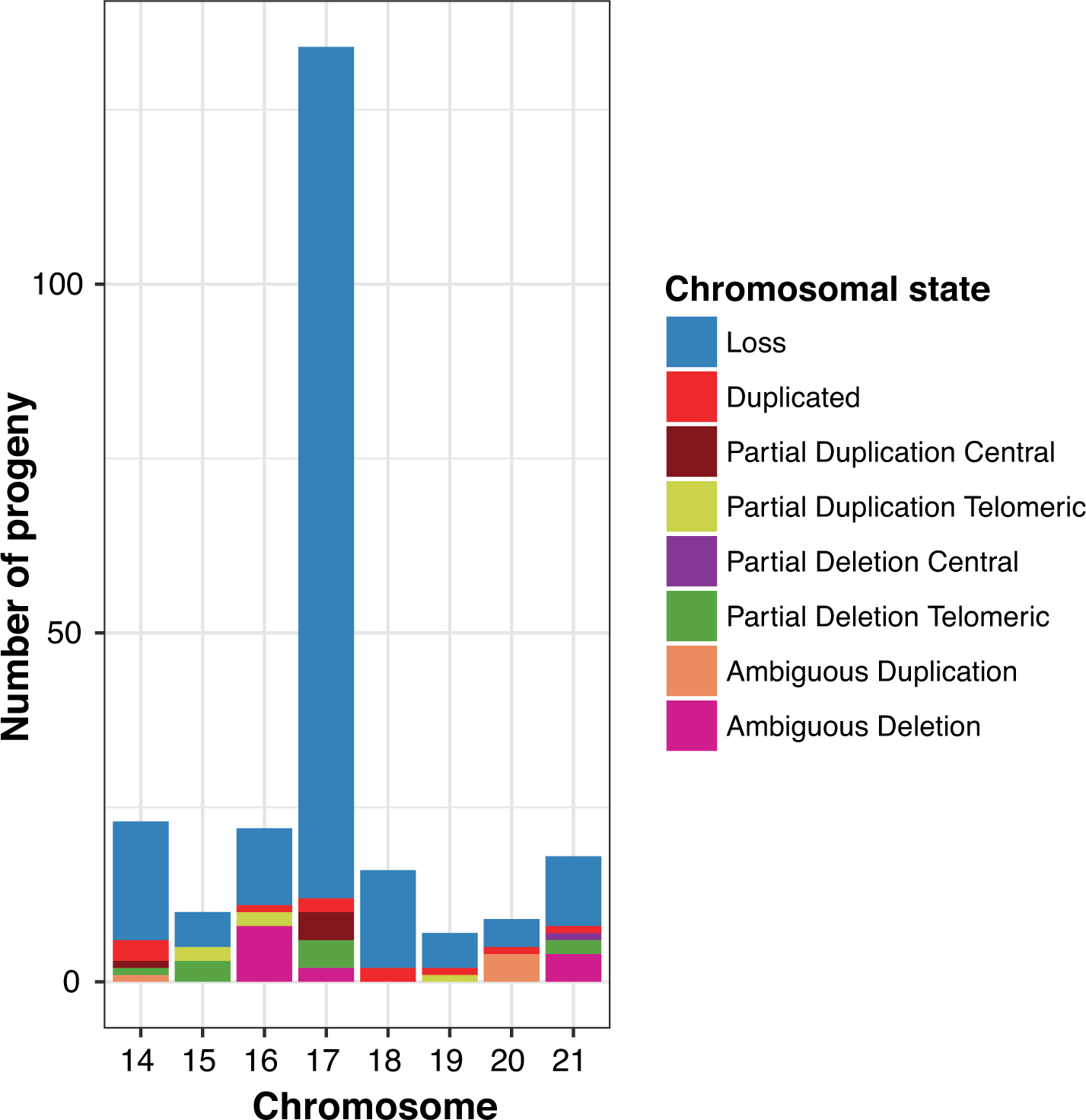
Identification of partial chromosome losses or duplications in cross 1E4x1A5. A summary of all the chromosome number and length polymorphisms in the progeny of cross 1E4x1A5, as well as the location where the length polymorphism occurred. Most of the rearrangements were ambiguous (19), 15 were located towards the ends of chromosomes and 6 rearrangements occurred in the central region of the chromosomes.

### Correlation of chromosomal features with the fidelity of transmission

During meiosis, chromosomes pair prior to recombination and therefore length similarity could play a role in homolog identification and enable chromosomes to pair and recombine. However, we found no correlation between the length similarity of the parent chromosomes and the fidelity with which chromosomes were inherited (Figure 6A). In general accessory chromosomes were more unstable than core chromosomes. Interesting exceptions were a disomic core chromosome 13 (length difference 5% between the parents) and a disomic core chromosome 5 (length difference of 8.4% between the parents). The rate of disomy for these core chromosomes was 1.6%. We found no significant correlation between the recombination rate and chromosome transmission fidelity (Figure 6B). However, in cross 1A5x1E4, most of the chromosome losses and disomies occurred in accessory chromosomes with a low recombination rate (Figure 6B). Next, we analyzed sequence similarities between parental chromosomes and correlated this with the chromosome transmission fidelity. For this, we compared whole chromosome sequences and calculated the percentage of syntenic regions between homologous chromosomes. The accessory chromosomes in the parents for both crosses had a much lower synteny than the core chromosomes and had substantially lower transmission fidelity (Figure 6C). Accessory chromosomes had overall a higher content in repetitive elements, which is similarly correlated with low transmission fidelity (Figure 6D).

**Figure 6:**
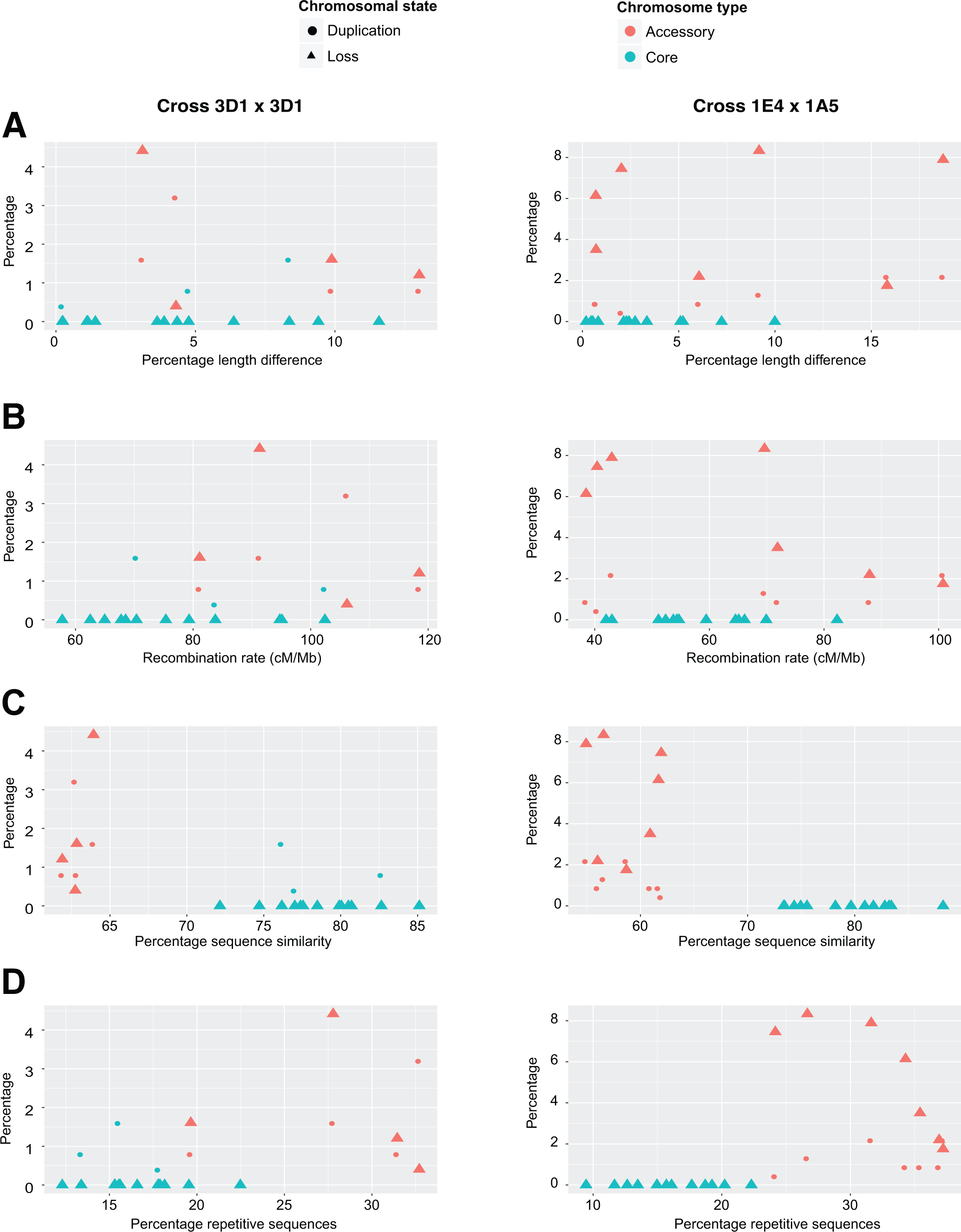
Correlations between chromosome length similarity, recombination rate, percent sequence similarity, fraction of repetitive sequences and the inheritance of chromosomes. Complete and partial chromosome losses and duplications were correlated with length similarity (A), recombination rate (B), sequence similarity (C) and repeat content (D) of the parental chromosomes. Correlations are shown separately for crosses 3D1x3D7 and 1E4x1A5.

### Association between accessory chromosomes and phenotypic traits

We analyzed whether the chromosome states in progeny were correlated with variation in phenotypic traits. For this, we considered first only two chromosome states: normal (haploid) or abnormal (any loss, duplication or rearrangements). We tested for an association with phenotypic traits using two-tailed *t*-tests (multiple testing significance threshold at *p* < 0.002). We first tested for associations with virulence on two wheat cultivars (Runal and Titlis) using data from a previous study (Stewart and McDonald, 2014; Stewart et al., 2017). Progeny from cross 3D1x3D7 with a normal chromosome 17 had a higher pycnidia count on the cultivar Runal than isolates with an abnormal chromosome 17 (*p* = 0.0019; Figure 7A). Isolates missing chromosome 17 had a lower pycnidia count than isolates that were disomic for chromosome 17. On cultivar Titlis, progeny from cross 3D1x3D7 with a normal chromosome 18 had significantly darker pycnidia (*p* = 0.0018). Progeny with an abnormal chromosome 19 had a marginally higher percent leaf area covered by lesions (PLACL; *p* = 0.0024). For progeny from cross 1E4x1A5, we found a correlation of the PLACL produced on Titlis with chromosome 21 (*p* = 0.00002; Figure 7B) and with chromosomes 15, 19 and 20 (*p* < 0.002). Isolates with a partially deleted or lost chromosome 21 had a higher PLACL. For progeny of cross 1E4x1A5, we found that isolates with an abnormal chromosome 20 showed higher PLACL on Runal. We found no significant correlations for phenotypes related to growth, fungicide resistance or temperature sensitivity.

**Figure 7:**
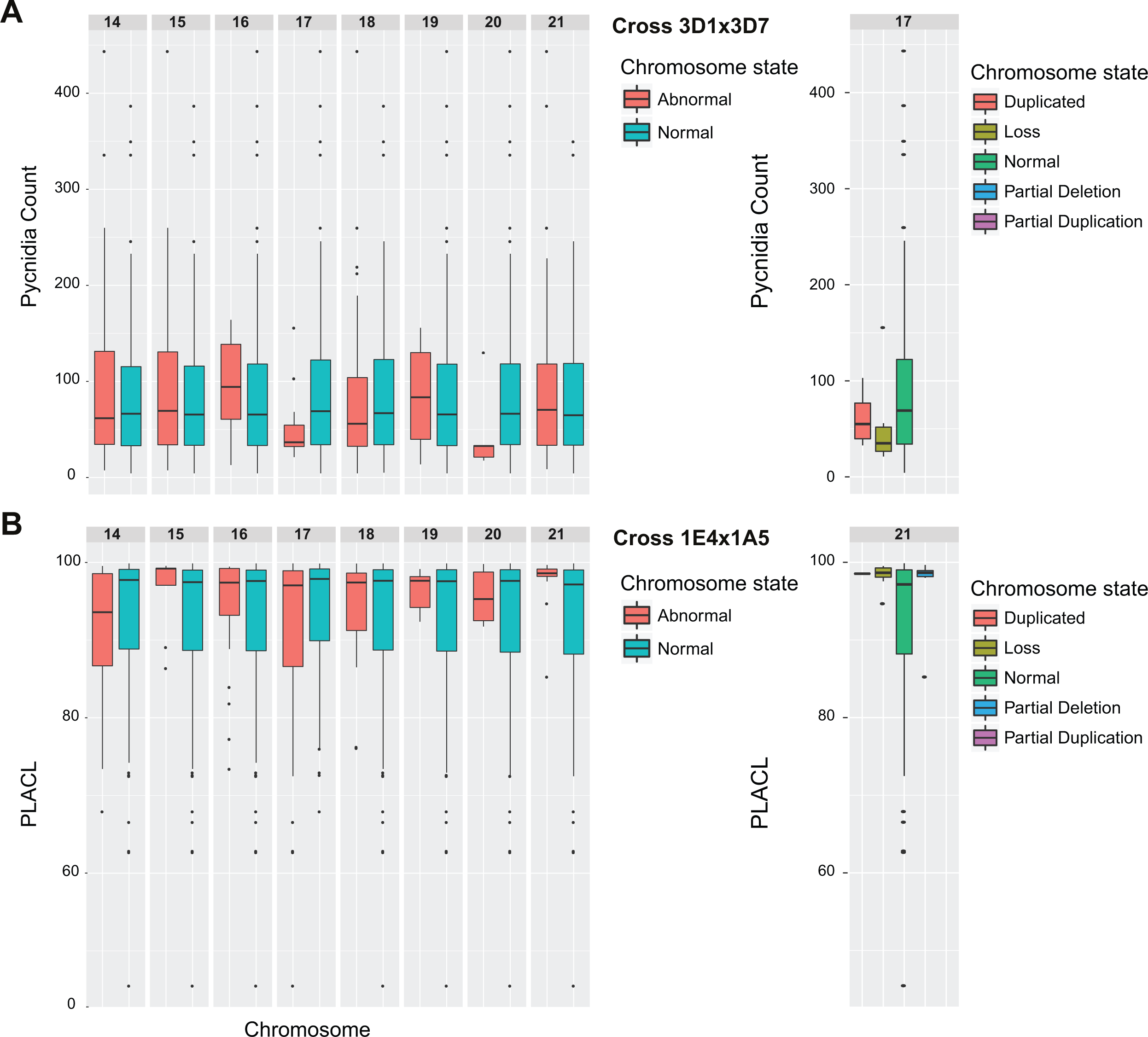
Association between accessory chromosomes and phenotypes. Accessory chromosome states, normal or abnormal (duplicated, lost, partially duplicated or partially lost), were compared to virulence traits using a two-sample t-test (multiple testing correction threshold of *p* < 0.002). (A) In the progeny of 3D1x3D7 isolates with a normal chromosome 17 had a significantly higher pycnidia count on the wheat cultivar Runal that isolates with a duplicated or lost chromosome (*p* = 0.0019). (B) In cross 1E4x1A5, isolates with a lost or partially deleted chromosome 21 had a higher percent leaf areas covered by lesions (PLACL) on Titlis than isolates with a normal chromosome 21 (*p* = 0.000024).

### Correlation between disomy and chromosomal rearrangements

We analyzed whether rates of disomy were correlated with rates of rearrangements. Non-disjunction results in the loss of a chromosome in one progeny and a chromosome gain in the corresponding twin spore from the same ascus. Core chromosomes generally showed only very rare cases of disomy or rearrangements (Figure 8). Accessory chromosome 14 was more frequently disomic and rearranged in progeny from cross 1A4x1E5. Chromosome 15 underwent partial duplications and deletions, but we found no evidence for non-disjunction. Chromosome 16 was both frequently rearranged (4.4%) and disomic (0.4%) among the progeny in 1E4x1A5. In cross 3D1x3D7, chromosome 17 was disomic in 1.6% of the progeny, while in cross 1E4x1A5 chromosome 17 was more rarely disomic (0.9%). Chromosome 17 showed even stronger differences in rearrangements among crosses (4.4% versus 0.0%). Chromosome 19 was both more likely to undergo rearrangements and to be inherited as a disomic chromosome in cross 3D1x3D7. In contrast, chromosome 21 was both more likely to be rearranged and to be inherited in a disomic state in cross 1E4x1A5.

**Figure 8:**
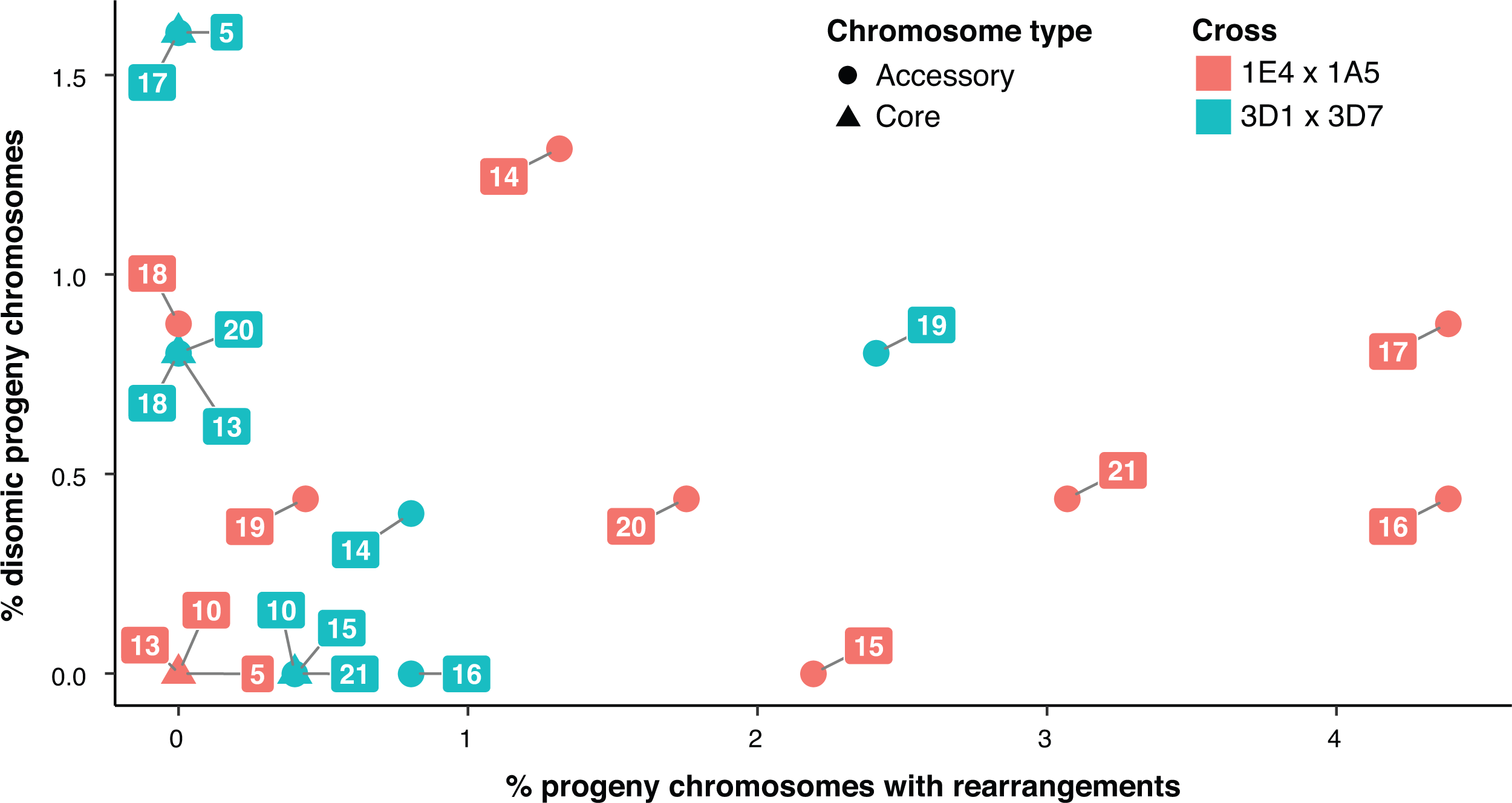
Correlation between number of disomic progeny and chromosomal rearrangements Circles and triangles represent accessory chromosomes and core chromosomes, respectively. Chromosomes from cross 3Dx3D7 are represented in blue, and chromosomes from cross 1E4x1A5 are in red.

## Discussion

We used RAD-seq data generated for several hundred progeny from two crosses of *Z. tritici* to identify aberrations in chromosomal transmission through meiosis. We found extensive chromosome number and length variation among the progeny in both crosses. The rates of disomy and rearrangements differed greatly between chromosomes and crosses. Nearly all aberrant chromosomal transmission events affected accessory chromosomes with the rare exception of core chromosome disomies. Several accessory chromosomes showed strongly distorted segregation frequencies.

Chromosome number polymorphism in *Z. tritici* has previously been linked to errors occurring during meiosis (Croll et al., 2013; Wittenberg et al., 2009). In our study, we generated a substantially more dense marker coverage using the Illumina-based sequencing technique RAD-seq and were able to screen more isolates (477 isolates compared to 144 and 216 isolates, respectively (Croll et al., 2013; Wittenberg et al., 2009)). Because RAD-seq generated a high coverage of ~100 bp sequences at defined restriction sites, we could precisely map sequences to chromosomal positions without having to rely on genetic map constructions. Physical marker positions are particularly important for analyzing accessory chromosomes of *Z. tritici*, because of their very low rates of recombination (Croll et al. 2015). In contrast to previous studies, our use of RAD-seq markers allowed us to directly detect duplicated chromosomal segments by analyzing variations in sequencing coverage.

Our analysis revealed that all eight accessory chromosomes underwent chromosome loss during meiosis. The rate of chromosomal loss depended on the chromosome and varied between the crosses. This confirms the findings of Croll et al. 2013, except that a loss of chromosome 15 had not previously been detected. We found that 5 progeny (2.1%) had lost this chromosome. No isolate was found lacking a core chromosome despite screening 477 progeny. This showed that all 13 core chromosomes are likely encoding essential functions for the growth and survival of the fungus. Chromosome loss most likely occurred as a result of errors during chromosome segregation, specifically non-disjunction of sister chromatids during either meiosis I or II. In accordance with previous studies, we found that the loss of accessory chromosomes during meiosis is common. In natural populations, this may lead to the complete loss of an accessory chromosome in the absence of counteracting mechanisms that maintain these chromosomes.

Wittenberg et al. (2009) proposed that distorted segregation of accessory chromosomes could serve as a mechanism to prevent their complete loss from a population. Chromosomes present in only one parent are expected to segregate into 50% of the daughter cells. However, we found that in cross 3D1x3D7 chromosomes 14, 15, 18 and 21 from parent 3D1 were significantly over-represented in the progeny. The transmission advantage resulting from unequal segregation is referred to as “meiotic drive” and is frequently associated with accessory or B chromosomes (Jones, 1991). Distorted segregation was not universal, for example chromosome 17 in cross 1E4x1A5 segregated normally. The distorted segregation pattern in cross 3D1x3D7 could be explained if parent 3D1 already had disomic accessory chromosomes. But our coverage analysis did not detect disomic chromosomes in any of the parents. The over-representation of progeny carrying a specific accessory chromosome could be due to selection favoring progeny carrying this chromosome or a meiotic drive mechanism. A possible explanation for the distorted inheritance of the accessory chromosomes is that some *Z. tritici* chromosomes may have “sticky” centromeres similar to those found in rye B chromosomes where the transmission at higher than Mendelian frequencies was explained by the presence of particular centromeres that ensure that B chromosomes migrate to the generative pole that will be transmitted to the next generation of plants (Banaei-Moghaddam et al., 2012). In order to distinguish between the exact mechanisms leading to distorted segregation, all meiotic products from individual tetrads would have to be analyzed. However, experimental limitations in the generation of large numbers of individual tetrads prevent us from making more detailed investigations using tetrad analysis.

We found that an average of 5.9% of the progeny isolates were disomic for one or more chromosomes. This number is similar to what was found for *Saccharomyces cerevisiae,* where 8% of the lab strains were estimated to be aneuploid (Hughes et al., 2000). Disomy is generated when chromosomes undergo non-disjunction during meiosis, resulting in one daughter cell with two copies of a chromosome and one daughter cell with no copies of that chromosome (Figure 3). Therefore, for each disomic offspring, we expect a corresponding offspring that is missing the same chromosome. As expected, we found that chromosomal loss was often accompanied by disomy. However, contrary to expectation there was no symmetry in the loss and disomy rates. For example, despite finding many progeny lacking chromosome 15, no isolate disomic for chromosome 15 was recovered. The rates of non-disjunction also differ between chromosomes and between crosses, suggesting that the loss or disomy of specific chromosomes may be counter selected. In addition, chromosomes differ in their composition of repetitive elements. Repetitive elements are likely to play an important role by influencing the likelihood of faithful disjunction. We also found that non-disjunction was happening during both meiosis I and II. We found heterozygous disomic chromosomes, which were created as a result of non-disjunction in meiosis I. Heterozygous disomic chromosomes were most frequent in cross 3D1x3D7. In cross1E4x1A5, homozygous disomy resulting from non-disjunction in meiosis II occurred more frequently. Aneuploidy plays an important role in adaptive evolution of fungal pathogens. In human pathogens, aneuploidy is often associated with drug resistance (Hu et al., 2008; Selmecki et al., 2010). Over 50% of the fluconozole-resistant strains isolated from patients had whole or partial chromosome duplications (Selmecki et al., 2006). Correlations between disomic states and phenotypic traits in *Z. tritici* suggests that selection could also be affecting rates of disomy, albeit with less drastic impacts than in human pathogens selected for drug resistance.

Aneuploidy typically causes a dosage imbalance, which could explain why accessory chromosome aneuploidies are tolerated more frequently than core chromosome aberrations. Alternatively, gene expression or dosage compensation could have evolved on frequently disomic chromosomes, which may explain the tolerance for additional copies of certain chromosomes, but not others (Torres et al., 2008). Chromosomes that have a higher rate of disomy could have shorter or non-functional telomeres. Telomere defects were found to explain mitotic instability in human mammary epithelial cells (Pampalona et al., 2010). Chromosomes with shorter telomeres are more likely to undergo non-disjunction. Furthermore, chromosomes with higher degrees of synteny are more likely to pair correctly, resulting in fewer non-disjunction events. We found indications that sequence similarity in the parent chromosomes indeed leads to higher fidelity of chromosomal inheritance.

Homologous chromosomes of *Z. tritici* segregate significant structural variation in populations, differ in repeat and gene content, chromosomal length, recombination rate, telomere and centromere composition (Croll et al., 2013, 2015; Plissonneau et al., 2016; Schotanus et al., 2015). Synteny breakpoints are commonly associated with repetitive sequences or transposable element clusters that can misalign during recombination, thereby generating length polymorphism. Such a mechanism was thought to generate a novel chromosome 17 in the progeny of cross 1A5x1E4 (Croll et al., 2013). In our study, we found no correlation between length similarity and recombination rate of the parent chromosomes, and the fidelity of chromosome inheritance. However, chromosomes with higher synteny between the parents and fewer repeats were transmitted more faithfully.

Selection favoring the presence or absence of specific accessory chromosomes would require that accessory chromosomes directly or indirectly influence phenotypic traits. However, accessory chromosomes carry few genes and none are thought to perform a specific function during the life cycle of the fungus (Goodwin et al., 2011). Interestingly, we found a correlation between the presence of chromosome 15, 18 and 21 and higher levels of virulence in cross 3D1x3D7 (Stewart et al., 2017). In addition, we found a correlation between the presence of a normal chromosome 17 and an abnormal chromosome 19 and higher levels of pycnidia and PLACL, respectively. In cross 1E4x1A5, abnormal chromosome 21 was associated with higher levels of PLACL. Even though the individual effect sizes were relatively small, the differences in the expression of virulence traits may be significant under natural conditions. The production of lesions (expressed as PLACL) and pycnidia counts can increase the survival and reproductive potential of the pathogen in the field. If isolates harboring specific accessory chromosomes gain a fitness advantage, then accessory chromosomes may be maintained in the species pool by a selection-drift balance.

Most chromosome rearrangements are thought to be deleterious and therefore counterselected. The ability of *Z. tritici* to tolerate a large number of disomies and chromosomal rearrangements makes this species an excellent model for detailed analyses of rearrangements and non-disjunction events. Despite the fact that the meiotic machinery is highly conserved, the strength of selection against erroneous chromosomal transmission can differ widely among species. Relaxed selection on chromosomal transmission can lead to highly polymorphic chromosomal sets observed in some eukaryotic pathogens. Determining the trade-offs involved in maintaining chromosomal integrity and generating chromosomal polymorphism will elucidate how selection operates to maintain the fidelity of meiotic processes.

## Acknowledgements

We thank Marcello Zala for providing access to progeny collections and helpful discussions. CP was supported by an INRA Young Scientist grant. SF and BAM are supported by the Swiss National Science Foundation (grant 31003A_155955). DC is supported by the Swiss National Science Foundation (grant 31003A_173265).

## Supplementary Figures and Tables

**Supplementary Table S1: Phenotypic trait data available for progeny.** PLACL: Percent leaf area covered by lesions. MC: Mean centred

**Supplementary figure 1: The normalized coverage distribution of progeny from cross 3D1x3D7 (A) and 1E4x1A5 (B).** Both normalized coverage distributions indicate the cutoffs used to detect a whole chromosome loss (ratio < 0.3), partial deletion (ratio 0.3-0.7), regular transmission (0.7-1.3), partial duplication (1.3-1.7) and whole chromosome duplication (>1.7). The normalized coverage was calculated by determining the coverage for each chromosome and comparing this to the median coverage of all chromosomes for that isolate

**Supplementary figure 2: Location of partial chromosome losses or duplications.** A summary of all the chromosome number and length polymorphism in the progeny, as well as the location where the length polymorphism occurred (cross 3D1x3D7).

**Supplementary figure 3: The coverage distribution on chromosome 10.** Isolates 3D1 and 3D7 are parental isolates. Progeny 89.1 displays a putative partial duplication of chromosome 10. Red and blue dotted lines show the genome-wide and chromosome wide median coverage, respectively.

